# Acinetobacter enrichment shapes composition and function of the bacterial microbiota of field-grown tomato plants

**DOI:** 10.1101/2025.07.14.664720

**Authors:** Senga Robertson, Alexandros Mosca, Saira Ashraf, Aileen Corral, Rodrigo Alegria Terrazas, Catherine Arnton, Peter Thorpe, Jenny Morris, Pete E Hedley, Giulia Babbi, Castrense Savojardo, Pier Luigi Martelli, Frederik Duus Møller, Hanne Nørgaard Nielsen, Pimlapas Leekitcharoenphon, Frank M. Aarestrup, Rashi Halder, Cedric C. Laczny, Paul Wilmes, Laura Pietrantonio, Pardo Di Cillo, Vittoria Catara, James Abbott, Davide Bulgarelli

## Abstract

Tomato is a staple crop and an excellent model to study host-microbiota interactions in the plant food chain. In this study we describe a ‘lab-in-the-field’ approach to investigate the microbiota of field-grown tomato plants. High-throughput amplicon sequencing revealed a three-microhabitat partition, phyllosphere, rhizosphere and root interior, differentiating host-associated communities from the environmental microbiota. An individual bacterium, classified as *Acinetobacter sp*., emerged as a dominant member of the microbiota at the plant-soil continuum. To gain insights into the functional significance of this enrichment, we subjected rhizosphere specimens to shotgun metagenomics. Similar to the amplicon sequencing survey, a ‘microhabitat effect’ defined by a set of rhizosphere-enriched functions was identified. Mobilisation of mineral nutrients, as well as adaptation to salinity and polymicrobial communities, including antimicrobial resistance genes (ARGs), emerged as a functional requirement sustaining metagenomic diversification. A metagenome-assembled genome (MAG) representative of *Acinetobacter calcoaceticus* was retrieved and metagenomic reads associated to this species identified a functional specialisation for plant-growth promotion traits (PGPTs), such as phosphate solubilization, siderophores production and reactive oxygen species detoxification, which were similarly represented in across tomato genotype-independent fashion. Our results revealed that the enrichment of a beneficial bacterium capable of alleviating plant’s abiotic stresses appears decoupled from ARGs facilitating microbiota persistence at the root-soil interface.

**IMPORTANCE:** Tomatoes are at centre-stage in global food security due to their high nutritional value, widespread cultivation, and versatility. Tomatoes provide essential vitamins and minerals, contribute to diverse diets, and support farmer livelihoods, making them a cornerstone of sustainable food systems. Beyond direct dietary benefits, the intricate relationship between tomatoes, their associated microbiota, and ARG is increasingly recognised. Tomato plants host diverse microbial communities in association with their organs, which influence plant health and productivity. Crop management impacts on composition and function of these communities, contributing to the prevalence of ARG in the soil and on the plants themselves. These genes can potentially transfer to human pathogens, posing a food safety and public health risk. Understanding these complex interactions is critical for developing sustainable agricultural practices capable of mitigating the impact of climatic modifications and the global threat of antimicrobial resistance.

## INTRODUCTION

Plants host diverse microbial communities in association with their organs, collectively referred to as the plant microbiota. Similar to the microbiota inhabiting vertebrates, the plant microbiota modulates the growth, development and health of its host (1). For instance, the microbiota at the root-soil continuum controls mineral biogeochemical cycles which, in turn, determines food provision as well as the environmental footprint of crop production (2). The plant microbiota does not represent a random assembly from the environment. Rather, a series of deterministic checkpoints define composition and function of the microbial assemblages thriving in association with plants (3). Consequently, the characterisation of the plant microbiota gained centre-stage as a first step towards the identification and development of “probiotics” for plants (4).

In recent years, plant-associated microbes have emerged as a neglected reservoir for antimicrobial resistance (AMR) mechanisms (5). Studies conducted with vascular plants and mosses identified antimicrobial resistance genes (ARGs) as microbiota determinants for host colonisation (6, 7). As ARGs are embedded in the microbiota at the plant-soil continuum (8), it is critical to define their dynamics across plant species to inform strategies aimed at improving ecosystem functioning and One Health (9).

Cultivated tomato (*Solanum lycopersicum*) is a staple crop (10) and an excellent experimental model for dissecting the genetic bases of plant-environment interactions (11). As tomato production is threatened by climatic modifications on a global scale (12), it is not surprising that innovative strategies to strengthen environmental resilience have been proposed, including an example of *de novo* domestication, whereby genome editing has been used to “convert” a wild relative into a of cultivated form (13). Among those strategies, the microbiota has gained centre-stage in both basic science and translational application as an untapped resource for climate change adaptation in crops (14), including tomato. For instance, the availability of tomato genetic resources enabled scientists to precisely identify host genetic determinants of the microbiota (15). Likewise, the threat posed by devastating soil-borne diseases has sparked efforts to identify a protecting microbiota capable of fending-off pathogens (16, 17). Despite the growing scientific interest into the tomato microbiota, available information remains focussed on taxonomic investigations (18). Predictive approaches to infer function from community composition have been applied (19–21), yet examples of characterisation of the tomato microbiota functional potential from soil-grown plants remains limited (22).

In this study, we embarked on a lab-in-the field approach, whereby sampling actual production sites in the Mediterranean environment we determined a “plant footprint” on the composition and function of the tomato bacterial microbiota which emerged sufficiently stable between the tested genotypes Motivated by the discovery of an unusual dominance of an individual phylotype classified as *Actinetobacter sp*., we further investigated the implications for agriculture, including antimicrobial resistance gene dynamics for One Health.

## MATERIAL AND METHODS

### Soil, plant and environmental samples

Sampling was performed in the summer of 2019 on a tomato farm close to the town of Larino [41°48′N 14°55′E], in the South of Italy. The farm grows distinct, processing-type, tomato F1 commercial hybrids: Abbundo [HM Clause; hereafter, ‘Ab’] and SV5197TP [Bayer; hereafter ‘SV’]. Plants were maintained in distinct fields of soils displaying comparable chemical and physical composition [Table S1]. Individual plantlets were transplanted in May and sampling was performed mid-June at early flowering stage by uprooting individual plants [Fig. S1]. Inter-row soil was used as ‘unplanted control’ representing the ‘bulk soil’ microhabitat. Samples were sealed in plastic bags and transported refrigerated to the University of Dundee. Crop management was identical for both hybrids and no significant difference in yield were recorded between hybrids [Pardo DiCillo, personal communication].

### Sample preparation and DNA extraction

Upon arrival, individual plants were shaken vigorously to remove any excess soil and debris. The resulting specimen of roots and rhizosphere soil was transferred to a 50 ml sterile falcon tube containing 15 ml phosphate buffered saline solution (PBS). Rhizosphere was extracted by vortexing for 30 s, samples were then transferred to a second falcon tube containing 15 ml PBS and the process was repeated to maximise rhizosphere recovery. The roots were removed from the sample tube and stored in a third PBS containing tube on ice for downstream preparation the same day. The rhizosphere suspensions were combined and centrifuged at 1,500 g for 20 min to pellet the rhizosphere, the PBS supernatant was discarded, and the pellet flash frozen using liquid nitrogen. Roots were removed from PBS storage and placed in sterile mortars, flash frozen with liquid nitrogen and finely crushed using a sterile pestle. Powdered root material was transferred to a sterile 15 ml falcon tube and flash frozen in liquid nitrogen. Leaf material was gently washed with sterile water and subjected to the same fine crushing protocol as described for root samples.

Unplanted soil was processed in a similar way to rhizosphere samples, 2-3 g of bulk soil was transferred to a sterile 50 ml falcon tube containing 15 ml PBS and subjected to the same centrifugation, supernatant disposal and flash freezing process.

All processed unplanted, rhizosphere, root and leaf samples were stored at -70 °C until further analysis. Each sample was subjected to DNA extraction using the FastDNA SPIN Kit for soil (MP Biomedicals, Solon, USA) according to the manufacturer’s instructions.

Six replicated swabs obtained from non-contiguous areas of each farm machinery used prior tomato harvesting and including, two tractors, two water hoses, a cultivator, a herbicide dispenser and the harvester were subjected to DNA extraction using the PowerSoil Pro kit (Qiagen, Manchester, UK) according to the manufacturer’s instructions.

### Amplicon preparation and sequencing

For bacterial amplicon profiling, the 16S rRNA hypervariable V4 region of the small ribosomal subunit rRNA gene was subjected to targeted amplification using the primer pair 515F (5’-GTGCCAGCMGCCGCGTAA-3’) and 806R (5’-GGACTACHVGGGTWTCTAAT-3’). The forward primers incorporate the Illumina flow cell adapter sequence at the 5’ termini and the reverse primers include unique 12 bp indexes incorporated to allow for multiplexing (23).16S rRNA gene PCR amplification was conducted using the KAPA HiFi Hotstart PCR Kit (KAPA Biosystems, Wilmington, USA). Metagenomic DNA (50 ng per sample) was used and individual reactions consisted of 4 μl 5X KAPA HiFi Buffer, 1 μl 10 mg/ml Bovine Serum Albumin (BSA), 0.6 μl 10 mM dNTPs, 0.6 μl 10 μM Forward primer, 0.6 μl 10 μM reverse primer, 0.25 μl KAPA hiFi polymerase, 2 μl 5 μM pPNA chloroplast blocker, 2 μl 5 μM mPNA mitochondrial blocker(24). The reactions were amplified using the following programme: 94 °C (3 min), followed by 35 cycles of 98 °C (30 s), 50 °C (30 s), 72 °C (1 min) followed by a final, single step of 72 °C (10 min). For each sample/index combination, a no template control (NTC) was run containing no DNA.

PCRs were performed in triplicate and using at least 2 individual master mix plates equating to six replicates per sample to minimise amplification biases. PCR reactions were pooled in an index-wise manner and 5 μl of each sample run on 1.5% agarose gel alongside its corresponding NTC. Only samples with clear banding at around 450 bp and where there was no detectable amplification of its corresponding NTC were taken forward. Amplicons were purified using Agencourt AMPure XP beads (Beckman Coulter, Amerhsam, UK) at a ratio of 0.7 μl beads to 1 μl sample. Individually indexed samples were pooled together in an equimolar ratio, supplemented with 20% PhiX control library run as recommended at a final concentration of 4 pM on an Illumina MiSeq with 2 x 150 bp chemistry. Sequencing of farm machinery derived metagenomic DNA was performed as previously described (25). Briefly, ∼5ng of isolated DNA was used for PCR amplification using primer pair 341F (5’-CCTACGGGNGGCWGCAG-3’) and 785R (5’-GACTACHVGGGTATCTAATCC-3’). An overhang adapter sequence was added in both primers. A second PCR was performed with primers against overhang sequences containing 8bp combinatorial dual indexes using product from first PCR as template. The purified second PCR product was pooled in equimolar concentration and was sequenced on MiSeq using 2x300 bp paired end reads.

### Amplicon data processing

Sequencing reads quality was assessed using FastQC, then DADA2 and R was used to generate ASV-count matrices using the basic methodology as outlined in the DADA2 pipeline tutorial as previously described (26). Briefly, read filtering was conducted using the DADA2 paired FastqFilter method, trimming 10 bp of sequence from the 5’ end of each read using a truncQ parameter of 2 and the maximum number of “expected” errors in a read (maxEE) of 2. The majority of amplicon reads remaining after this process were of high quality and did not require 3’ end trimming. The DADA2::Learn_errors() method was run to determine the error model with a MAX CONSIST parameter of 20 whereby the error model will converge or be close to convergence within 20 rounds. The DADA2::DADA() method was then run with the error model to de-noise the readings using sample pooling. Reads were then merged setting a minimum overlap of 12 bp [actual overlap in the final amplicon pool 45-49bp] followed by the removal of chimeras using the consensus method. The dada2::assignTaxonomy() method was used using the RDP Naïve Bayesian Classifier to assign taxonomy with the SILVA database version 138 (27) imposing a minimum bootstrap confidence of 50. Following taxonomic assignment, the outputs were converted into ‘phyloseq objects’ (27) for downstream statistical analyses and figure generation. The phyloseq object was pruned *in silico* of ASV classified as chloroplast and mitochondria as well as matching a dataset of putative contaminants of the lab (28). Likewise, we discarded ASV for which we were unable to determine affiliation at phylum level. Next, we imposed a secondary quality filtering (29) based on ASV abundance, retaining in the final dataset only the those accruing 20 reads in at least 1% of the samples (i.e., an individual sample or more). Following pre-processing, samples with more than 5,000 reads were retained for downstream analysis. Rarefaction curves were computed with the R package vegan using a pre-compiled code (30) while ecological indices, i.e., alpha and beta diversity, were computed with phyloseq. Upon assessing data distribution with a Shapiro-Wilk test, univariate statistical analysis was performed using either analysis of variance or a Kruskal-Walls test setting an alpha level of 0.05. Permutational analyses of variance on distance matrices were computed using the function adonis of the package vegan (31). We used DESeq2 (32) to identify differential enrichment as previously described (26).

### Co-occurrence networks

Co-occurrence networks were generated using ASV data agglomerated at genus level. Briefly, networks were generated with sparCC (33) considering only genera with a relative abundance greater than 0.05% and occurring in at least 50% of the samples in the set. Cytoscape (34) was adopted to visualise and analyse networks. Both positive and negative correlations with an absolute value greater or equal 0.50 were plotted in networks but only positive correlations have been retained to analyse network modularization. The Cytoscape gLay plugin (35) was used to identify modules - i.e., groups of nodes more tightly connected among each other than with the rest of the network. The NetworkAnalyzer plugin (36) was used to identify central genera. Three different centrality measures, i.e., degree, betweenness and closeness, were computed for each node in a network (36).For each network, the distributions of node centralities were computed independently for each measure. Nodes with at least one measure higher than the 90^th^ percentile were retained as central.

### Metagenomic data preparation and sequencing

Raw DNA samples (70 μl) were purified using Agencourt AMPure XP Kit with a ratio of 1.8 μl beads per 1 μl sample. The samples were eluted in Buffer EB with a final volume of 40 μl and their concentration measured by Qubit. The samples were then fragmented using QIAGEN FX DNA Library Core Kit. 110 ng sample was fragmented in a total volume of 50 μl including 5 μl 10X FX Buffer and 10 μl FX Enzyme Mix. Fragmentation was performed in a thermocycler using the following program: 4 °C for 1 min, 32 °C for 6 mins (varies with amount of DNA), 65 °C for 30 min, and finally holding sample at 4 °C. For samples with less than 110 ng of DNA, the FX reaction mix was made to a total volume of 60 μl with 6 μl 10X FX Buffer and 12 μl FX Enzyme Mix. The fragmentation time at 32 °C was also varied according to the amount of DNA as follows: 60-100 ng DNA – 7 min; 40-60 ng DNA – 8 min; 20-40 ng DNA – 9 min. Each sample had a unique adapter attached which allowed binding to the Illumina flow cell for sequencing purposes and the unique identification of the samples in the library. The QIAseq UDI Y-Adapter Kit was used and 5 μl of a unique adapter was added per sample. Subsequently, 20 μL 5X Ligation Buffer, 10 μl DNA Ligase, and either 15 or 5 μl Nuclease-free water was added per sample depending on the volume of the fragmented sample, to make a total volume of 100 μl. The fragmented samples were then incubated at 20 °C for 15 min in a thermocycler with the heated lid off. Immediately afterwards, the samples were cleaned up using 0.8X (80 μl) Agencourt AMPure XP beads and eluted in 52.5 μl of Buffer EB. A second purification was performed using 1X (50 μl) Agencourt AMPure XP beads and elution in 26 μl Buffer EB. After adapter ligation and sample purification, PCR amplification was performed on the DNA samples. To 23.5 μl sample DNA, 25 μl QIAseq 2X HiFi PCR Master Mix and 1.5 μl QIAseq Primer Mix (10uM each) was added. PCR was then performed in a thermocycler using the following program: 98 °C for 2 min, followed by 10 cycles of 98 °C for 20 s, 60 °C for 30 s and 72 °C for 30 s, and finally 72 °C for 5 min before holding samples at 4 °C. After performing PCR, the samples were purified using 1X (50 μl) Agencourt AMPure XP beads and eluted in 25 μl Buffer EB. The samples were checked for amplification by gel electrophoresis. 5 μl of each sample was run on a 1.5% agarose gel. A fragmented pattern with the highest intensity centred at around 550 bp indicated successful amplification. The DNA samples were then quantified using the Qubit 4 Fluorometer. An equimolar library was prepared and submitted for sequencing purposes. Metagenomics libraries were sequenced on Illumina NextSeq 2000 instrument using a P3 kit cycles (360 Gb) according to manufacturer’s recommendations.

### Metagenomic data processing

Sequencing reads were quality assessed using FastQC and quality/adapter trimmed using TrimGalore (37), using a quality cutoff of 20, a minimum sequence length of 100 bp, and removing terminal N bases. Taxonomic classification of the sequence reads was performed using Kraken 2.0.9 (38) with custom Kraken database consisting of the Kraken archaea, bacteria, plasmid, viral, human, fungi, plant and protozoa sections. Host contamination was removed by alignment against the tomato cultivar Heinz 1706 (AEKE03000000) genome sequences, using BWA MEM (39), and non-aligning reads were extracted from the resulting BAM files using SAMtools (40). Metagenome assembly was conducted using MegaHit version 1.2.9 (41) with the “meta-large” preset. Metagenome-assembled genomes (MAGs) were created using the MegaHit-assembled contigs described above and MetaBat2 version 2.15 (42) to create contig bins representing single genomes. Contig bins were dereplicated using dRep version 3.2.0 (43) followed by decontamination with Magpurify version 2.1.2 (44). The resulting MAGs were assessed for completeness and contamination using checkM (45). Annotation of the MAGs was performed with bakta version 1.9 (46) before taxonomic classification was determined using the Genome Taxonomy Database Toolkit (GTDB-Tk) version 1.4.0 (47) with data version r95 To refine MAGs’ identification, the processes of assembly, binning, and taxonomic identification were iteratively refined in two steps, conducted separately for both genotypes. In the first step, metagenomic reads were assembled after removing plant-derived sequences. In the second step, the remaining reads —i.e., those not included in the previous assembly— were reassembled. Using BWA MEM and SAMtools, the *Acinetobacter calcoaceticus* MAG was mapped against metagenomic reads that had been previously filtered to remove host sequences and the assembled reads from the first assembly.

Metagenomic reads classified as *Acinetobacter calcoaceticus* by Kraken 2.0.9 were extracted and subsequently indexed and mapped against the metagenomic dataset using BWA, applying the default parameters for minimum seed length (19) and minimum alignment score (30). The aligned metagenomic reads were assessed using the ‘flagstat’ command of SAMtools for the read alignment statistics, considering the only primary mapped reads of *A. calcoaceticus* on the metagenomic dataset. To further validate the taxonomic classification, we performed an average nucleotide identity (48) comparison in Proksee (49) with the reference genomes of *A. calcoaceticus* [strain NCTC12983] and *A. baumanni* [strain ATCC 19606] obtained from NCBI genome . PGPg Finder (v.1.10) (50), based on the comprehensive PlaBase tool version 1.01 (51) was used with the ‘metafast_wf’ set in order to uncover the plant-growth promoting traits in the metagenomic sequences obtained from the remaining reads prior the second assembly analysis and in the metagenomics reads of *Acinetobacter calcoaceticus.* PGPg Finder outputs were converted into a phyloseq object for statistical analysis and figures generation [alpha, beta diversity and differential enrichment analysis, see above]. Individual samples and rarefied at 400,000 reads prior calculation.

The additional assembled contigs have been mapped across the 37 samples using the jgi_summarize_bam_contig_depths command from MetaBat2, in order to estimate contig coverage in each samples. The gene functions of the contigs were then annotated using eggNOG-mapper (v. 2.1.10) (52). Subsequently, the resulting data were considered to perform a functional enrichment analysis with DESeq2, comparing the functional profiles of the rhizosphere and soil separately for ‘Ab’ and ‘SV’ genotypes.

### Resistome reconstruction and visualisation

Metagenomics reads, quality checked and pruned of putative host contamination (see above), were mapped to whole genome bacterial database from NCBI and ResFinder (53) for ARG using KMA alignment (54) and default parameters. ARG’s abundance was identified as FPKM (fragments per kilobase per million fragments). This accounted for both sample-wise sequencing depth differences and size-dependent probability of observing a reference (55). FPKM was calculated by first normalizing for read depth by dividing the read counts with the total number of reads aligned to bacterial database in the sample and using a scaling factor of 1 million. Then, the read counts were normalized according to gene length by dividing them with the length of the gene in kilobases (56). For heatmap construction, genes log(FPKM) values were used to narrow the range and enhance heatmap readability. Using the "vegdist" function from the Vegan R package, a distance matrix was generated based on Bray-Curtis dissimilarity, which quantifies differences between samples based on FPKM values. Sample clustering was executed using the "pheatmap()" function from the pheatmap R package, employing complete linkage hierarchical clustering on the Bray-Curtis distance matrix.

The reconstructed Acinetobacter MAG was submitted to the public ResFinder server and the The Comprehensive Antibiotic Resistance Database - Resistance Gene Identifier (57) for ARG identification using default and “stringent” parameters, respectively.

### Acinetobacter genomes phylogenetic analysis

We retrieved sixty-five publicly available *Acinetobacter calcoaceticus* genomes and categorised them as ‘plants/soil’ or ‘other’ according to isolation source information available on NCBI. Phylogenetic tree construction and annotation was performed as previously described (58).

## RESULTS

### The bacterial microbiota of field-grown tomato plants is a “gated” community

We generated 173 bacterial 16S rRNA gene amplicon profiles of four microhabitats, i.e., unplanted soil, rhizosphere, roots and leaves, from two tomato F1 hybrids grown in adjacent fields in the south of Italy. Upon an *in silico* pruning of ASVs putatively representing host or environmental contamination and applying an abundance threshold, we retained 129 samples: number of samples unplanted soil *N*_unplanted_ = 14; *N*_rhizosphere_ = 51; *N*_roots_ = 49; *N*_leaves_ = 15. Those samples accrued for 940 ASVs and 3,747,178 sequencing reads. These ASVs were affiliated to 20 phyla, although five, namely Acidobacteriota, Actinobacteriota, Bacteroidota, Firmicutes and Proteobacteria represented over 99% of the retained reads in plant’s associated microhabitats [Fig. S2].

Rarefaction curves indicated that we reached a plateau-like pattern for the vast majority of our samples [Fig. S3], motivating us to inspect ASV richness and evenness across microhabitat. This revealed a distinct pattern for the tomato bacterial microbiota, manifested by a significant reduction in both metrics when comparing plant-associated microhabitats, i.e., leaves, roots and rhizosphere, with unplanted soil controls [Figure 1A, B; Kruskal-Wallis test, individual *P* values <0.05]. Likewise, a dissimilarity matrix computed on microbiota composition partitioned individual samples in a microhabitat-dependent manner [Figure 1C, *R*^2^ ‘Microhabitat’ = 0.52, permutational analysis of variance, *P* value < 0.001; 5,000 permutations].

**Figure 1:**
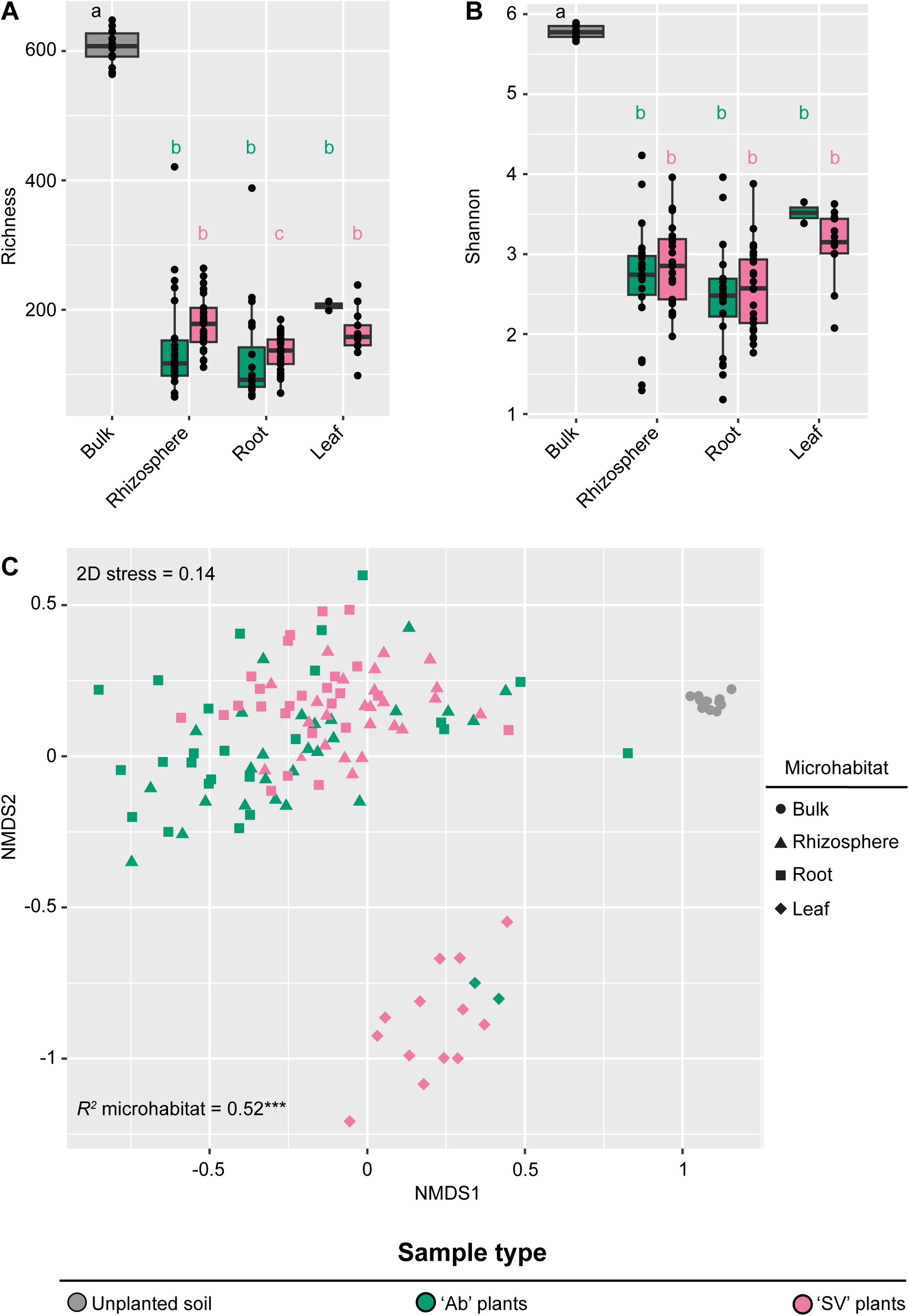
(**A**) Total number of observed ASVs and (**B**) Shannon’s diversity index, calculated for bulk and plant associated microhabitats. The upper and lower edges of the box plots represent the upper and lower quartiles, respectively. The bold line within the box denotes the median, individual shapes depict measurements of individual biological replicates/genotypes for a given microhabitat. Different letters denote statistically significant differences between microhabitat means by Kruskal–Wallis non-parametric analysis of variance followed by Dunn’s post-hoc test (Individual *P* values < 0.05; BH corrected). (**C**) Nonmetric multidimensional scaling computed using a Bray-Curtis dissimilarity matrix. Individual shapes depict replicates of indicated microhabitats, colour-coded according to sample type or genotype. The *R^2^* value depict the proportion of variation in distances explained by the factor microhabitats (adonis test, *P value* < 0.001; 5,000 permutations).

To gain further insights into the individual members of the microbiota underpinning the compositional diversification we took a two-pronged approach. First, we generate co-occurrence networks computed on bacterial genera relative abundance. Outputs of this investigation mirrored the previously identified pattern: while unplanted samples returned ten modules, plant-associated microhabitat were characterised by only three or four [Figure 2]. Likewise, network topology, defined by number of nodes and edges, emerged as more complex in unplanted soil compared to plant-associated microhabitats [Table S2]. A further discrimination emerged from the taxonomic inspection of these networks: in unplanted soil and leaves the group *Allorhizobium-Neorhizobium-Pararhizobium-Rhizobium* was more central than in roots and rhizosphere microhabitats. Conversely, belowground networks appeared supported by other Proteobacteria and Firmicutes [Table S2].

**Figure 2:**
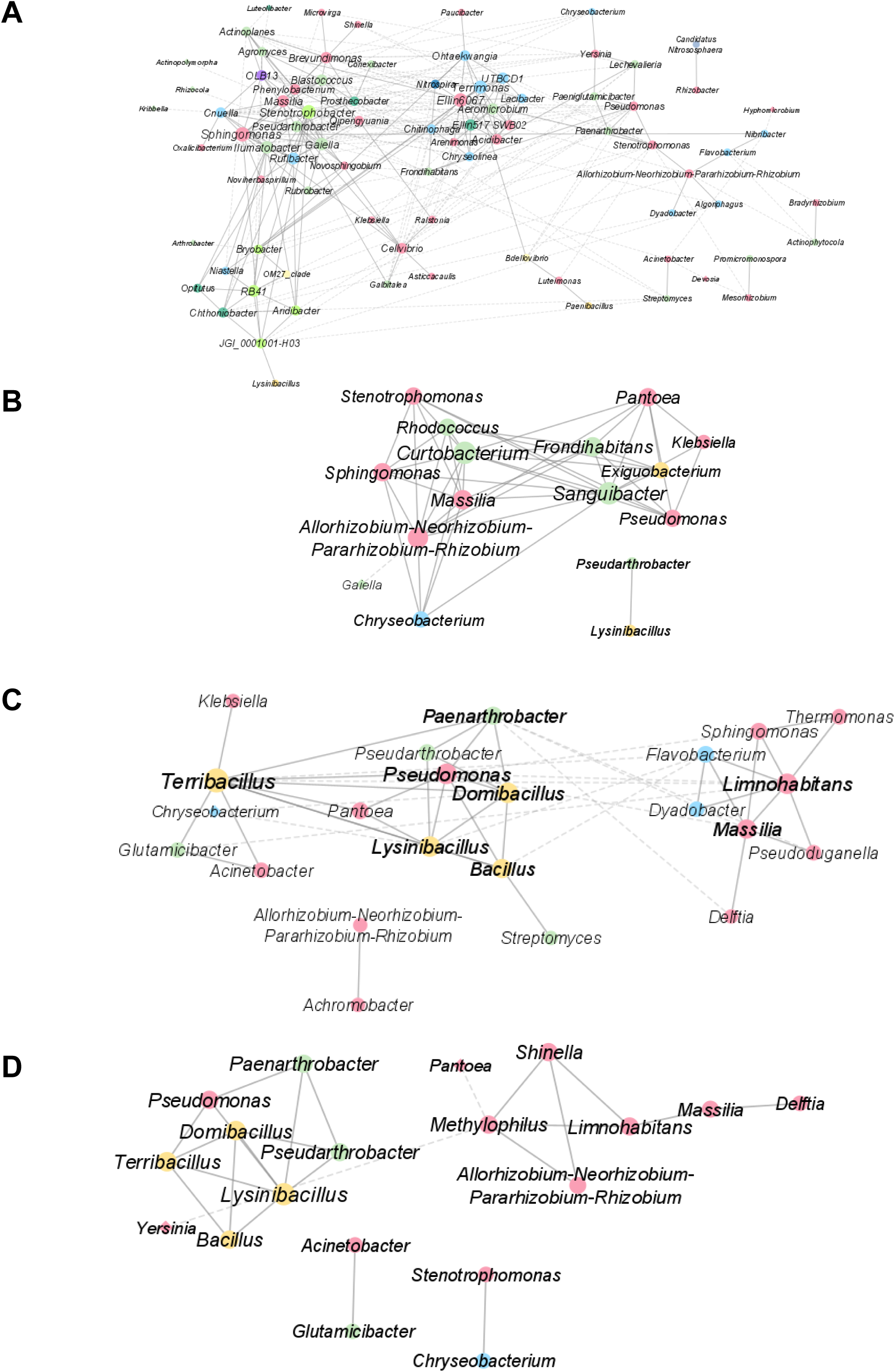
Correlation networks among bacterial genera for bulk (**A**), leaves (**B**), rhizosphere (**C**), and root (**D**) samples. Nodes are coloured following the bacterial phylum. Links depicted with solid lines and dashed lines represent positive and negative correlations, respectively. Node dimension is proportional to the node degree.

Next, we performed pair-wise comparisons between unplanted soil with root and rhizosphere specimens, respectively. We identified ASVs significantly enriched in and diagnostic for plant-associated microhabitats, although their numbers appeared orders of magnitude lower than what previously reported for the species (18) [Figure 3, Wald test, individual *P* values < 0.05; FDR corrected; Data Set S1 at https://zenodo.org/records/17580416]. Closer inspection revealed that an individual ASV, classified as *Acinetobacter sp.*, accrued for ∼30% of sequencing reads on average in both root and rhizosphere specimens, regardless of the genotype investigated [Fig. S4]. As a dominant Acinetobacter is likely to occupy niches which would otherwise be available to other members of the microbiota, this may explain, at least in part, the reduced number of observations compared with other studies.

**Figure 3:**
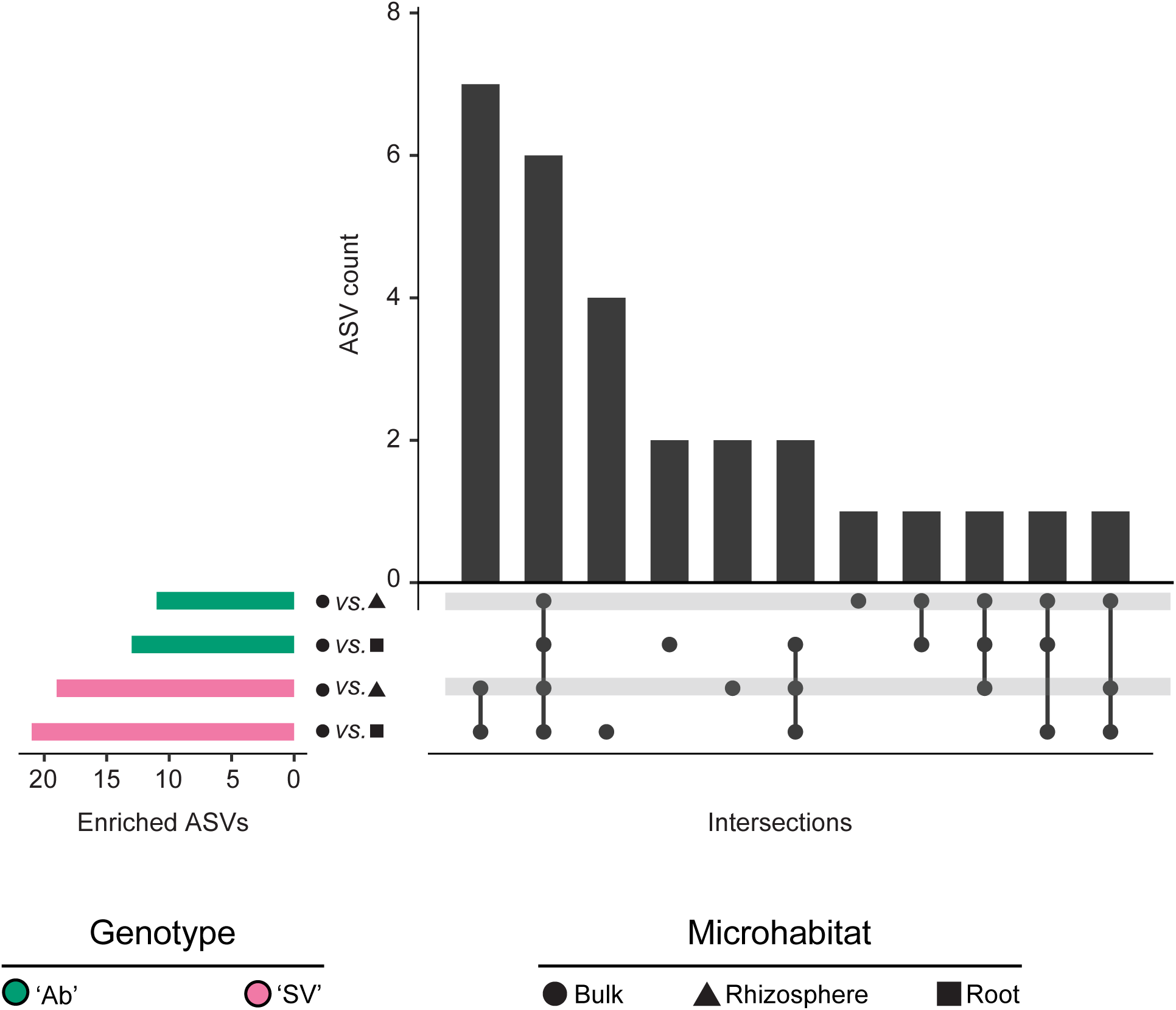
Horizontal blue bars denote the number of ASVs differentially enriched (Wald test, individual *P values* of <0.05, FDR corrected) between the bulk soil and root and rhizosphere microhabitats in the two tomato genotypes as recapitulated by the shape and colour scheme. Vertical bars depict the number of differentially enriched ASVs unique for or shared among two or more pairwise comparisons highlighted by the interconnected dots underneath the vertical bars.

This surprisingly high proportion of an individual bacterium motivated us to inspect microbiota composition of seven farm machineries as possible sources of an artificially derived *Acinetobacter* contamination. Although different amplification protocols prevented us to perform a deeper taxonomic comparison, the farm machinery microbiota resulted dominated by members of the phylum Actinobacteriota [mean reads proportion ∼39% across samples; Fig. S5]. Conversely, *Acinetobacter sp*. was detected in the same order of magnitude of bulk soil specimens [mean reads proportion ∼1.5% across samples; Fig. S5]. We therefore concluded observed Acinetobacter dominance represents a *bona fide* endogenous trait of the field-grown tomato microbiota.

### A three-pronged functional specialisation of the tomato rhizosphere bacterial microbiota

To gain insights into the functional significance of the bacterial microbiota for field-grown tomato plants, we reconstructed 37 bulk soil and rhizosphere metagenomes and classified ∼675 million paired-end reads. Taxonomic profiles of metagenomic samples were dominated by bacteria [> 637 million reads, ∼94% classified reads; Data Set S2 at https://zenodo.org/records/17580416] and mirrored the ones obtained from cognate amplicon samples and returned a limited, but significant, correlation between the two methodologies [Fig. S6; Spearman’s rank correlation rho = 0.17; P value = 0.0003]. This observation suggests that the footprint of host selection at the root-soil interface is sufficiently resistant to sequencing bias.

This motivated us to mine a repository of plant-growth promoting traits (59) to determine functional specialisations of the tomato rhizosphere microbiota. This allowed us to identify 2,642 individual microbial genes putatively representing plant growth-promoting traits. Inspection of gene distribution between microhabitats revealed an opposite trend compared to amplicon sequencing profiles. Irrespective of the plant genotype tested, the rhizosphere microhabitat emerged as richer and more diverse functional profiles than bulk soil controls [Figure 4A, B; Kruskal-Wallis test, individual *P* values <0.05] while functional gene abundance returned a clear microhabitat-dependent partition of individual samples [Figure 4C, *R*^2^ ‘Microhabitat’ = 0.68, adonis test, *P* value < 0.001; 5,000 permutations]. This apparent incongruence between microbial richness can be explained by the fact that plant-growth promoting traits are likely to be enriched in plant-associated samples compared to unplanted controls. Consistently, the rhizosphere enrichment of 15 functional categories out of 142 accrued for over half the sequencing reads and significantly discriminate between microhabitats [Figure 4D, Wald test, individual *P* values < 0.05; FDR corrected; Data Set S3 at https://zenodo.org/records/17580416]. These functional categories defined a three-pronged specialisation of the tomato microbiota: alongside genes required for the mobilisation and plant’s uptake of mineral nutrients, we identified genes putatively implicated in adaptation to abiotic stresses as well as modulation of inter-organismal relationships. Among the latter, we identified genes putatively implicated in antimicrobial resistance. We therefore reconstructed a resistome representative of the metagenomic dataset. We were able to identify 25 ARGs with sufficient confidence. Despite no clear microhabitat- or genotype-dependent pattern could be discerned, genes coding for beta-lactamase emerged as dominating the sequencing profiles [12 out of 25 identified genes; Figure 5].

**Figure 4:**
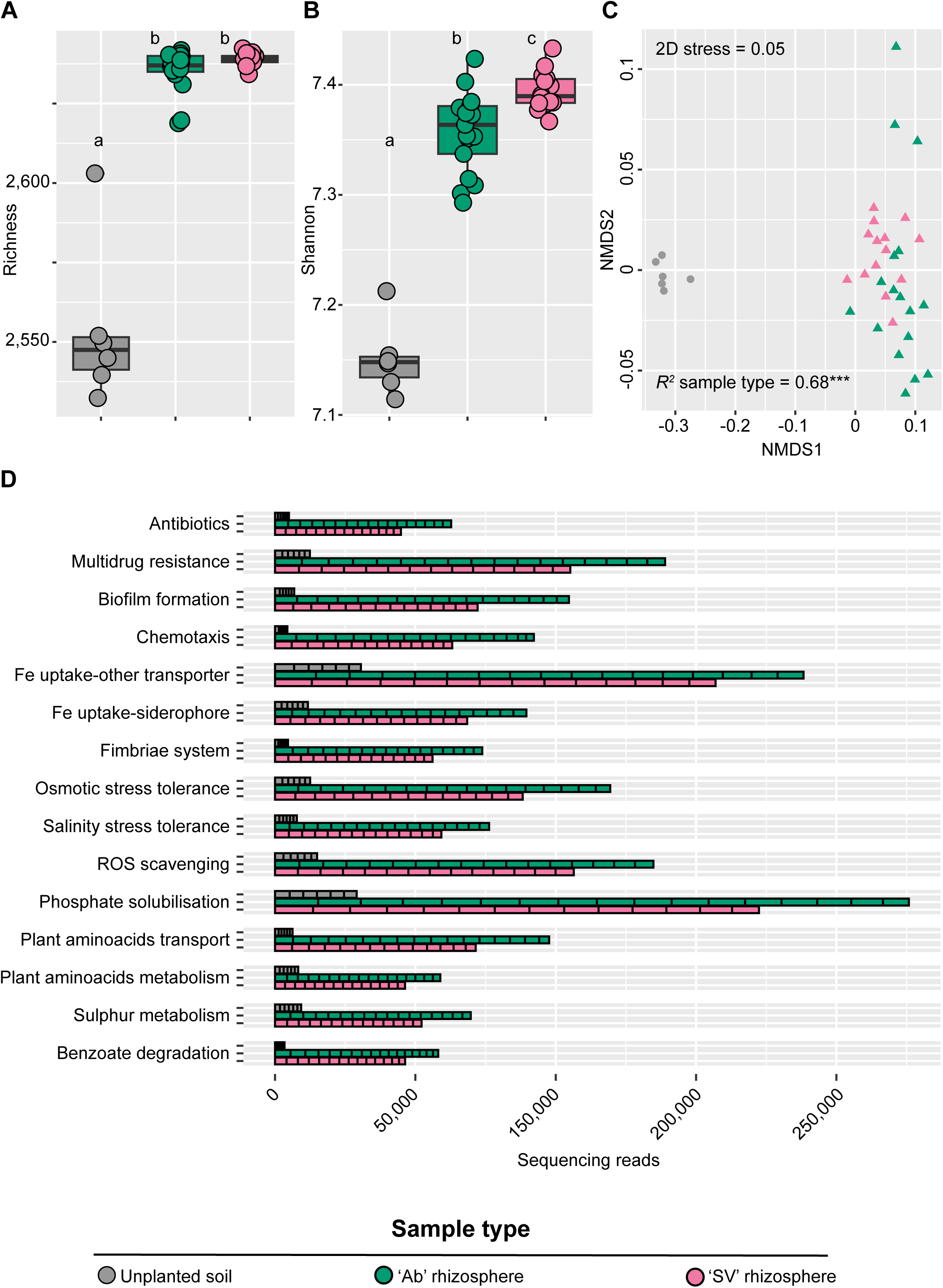
(**A**) Total number of observed functions and (**B**) Shannon’s diversity index calculated for bulk and rhizosphere microhabitats. The upper and lower edges of the box plots represent the upper and lower quartiles, respectively. The bold line within the box denotes the median, individual shapes depict measurements of individual biological replicates/genotypes for a given microhabitat. Different letters denote statistically significant differences between microhabitat means by Kruskal–Wallis non-parametric analysis of variance followed by Dunn’s post-hoc test (Individual *P* values < 0.05; BH corrected). (**C**) Nonmetric multidimensional scaling computed using a Bray-Curtis dissimilarity matrix on functional genes. Individual shapes depict replicates of indicated microhabitats, colour-coded according to sample type or genotype. The *R^2^*value depict the proportion of variation in distances explained by the factor microhabitat (adonis test, *P value* < 0.001; 5,000 permutations). (**D**) Cumulative abundance of genes representative of 15 most dominant functional categories enriched in and differentiating between rhizosphere and bulk soil in both genotypes enriched (Wald test, individual *P values* of <0.05, FDR corrected).

**Figure 5:**
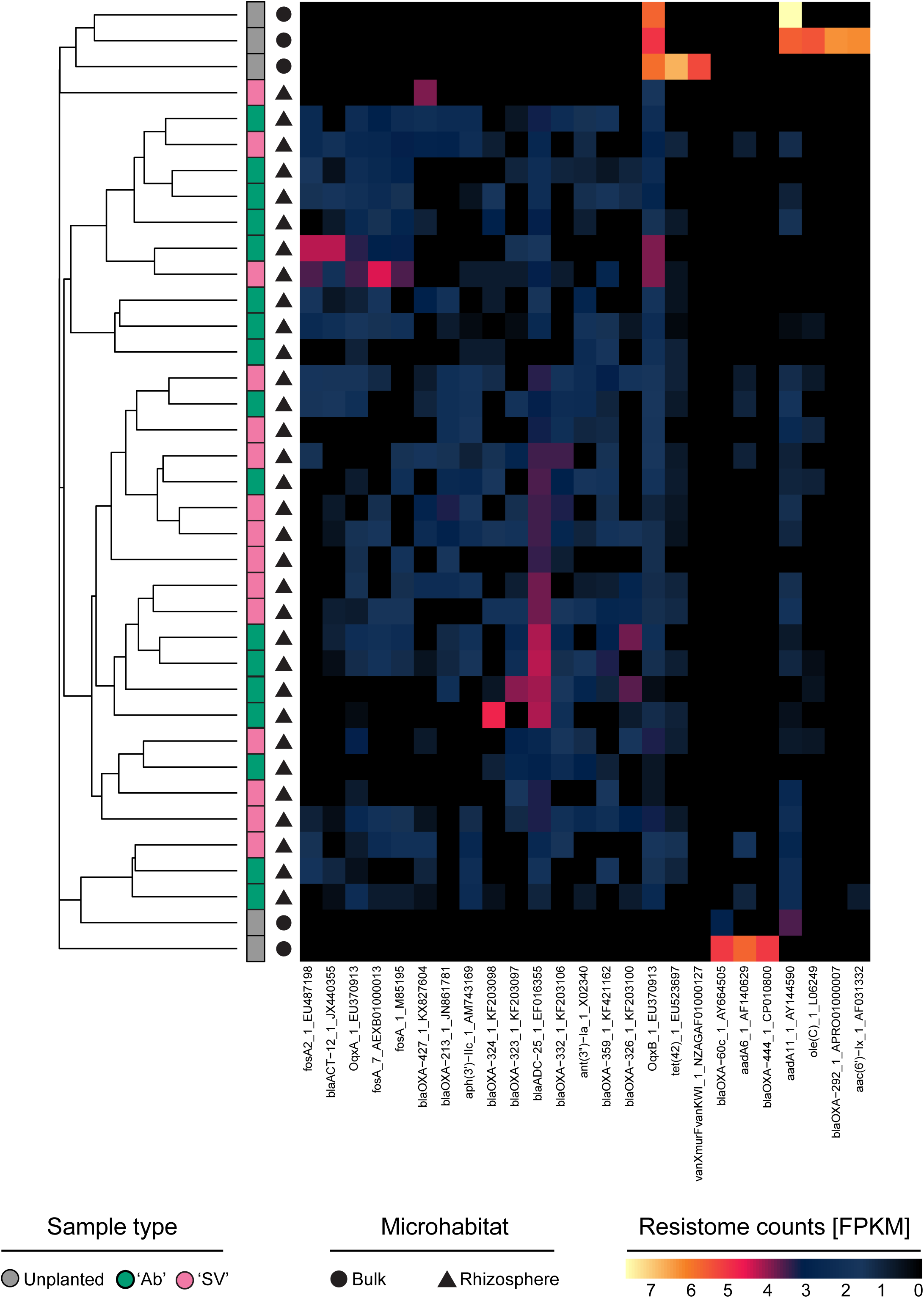
Abundance of individual ARGs, identified in the indicated microhabitat and tomato genotypes. Abbreviations: *fos*, fosfomycin resistance genes; *bla*, beta-lactamase; *Oqx*, efflux pumps, fluoroquinolone; *aph*, aminoglycoside 3’-phosphotransferase; *ant*, Aminoglycoside nucleotidyltransferase; *tet*, tetracycline resistance genes; *aadA*, aminoglycoside adenylyltransferase; *oleC*, oleandomycin; *aaC*, aminoglycoside acetyltransferase.

### Plant-growth promoting functions of *A. calcoaceticus* are stable between host genotypes

To further investigate the potential plant-growth promotion traits (PGPTs) of the dominant member of the bacterial communities, we attempted at reconstructing metagenome-assembled genomes phylogenetically related to *Acinetobacter sp*. First, we assembled over 1.5 Gbp metagenomic reads from the two tomato genotypes [Table S3] from which we were able to reconstruct an induvial *Acinetobacter sp*. MAG with completeness >70% and contamination <10%, taxonomically identified as *Acinetobacter calcoaceticus*. Comparison of the MAG sequences with the reference genome of *A. calcoaceticus* returned an average nucleotide identity of 94.99%, greater than the value of 86% obtained when the same analysis was performed against the reference genomes of *A. baumanni*. This motivated us to retrieve sixty-five publicly available *A. calcoaceticus* genomes to inspect phylogenetic relatedness and colonisation preference within the species. Although no clear partition according to ecological niche could be discerned, we identified two plants/soil isolates among the closest relatives of the identified MAG [Fig. S7]. This further supports the notion that the Acinetobacter dominance of the amplicon and metagenomic profiles may represent an endogenous colonisation of the field-grown tomato microbiota. Mapping analysis of the *A. calcoaceticus* MAG against the metagenomic reads, filtered to exclude plant host sequences and previously assembled reads, revealed a similar representativeness of this bacterial species across the two genotypes. In line with the ASV data, the prevalence of the mapped reads was in the rhizosphere samples, indicating a high representativeness of this MAG in specific ‘Ab’ rhizosphere samples, with the highest proportion of more than 10% among them [Table S4].

However, when we explored the abundance of PGPTs extracted from *A. calcoaceticus* reads, no host-genotype effect was identified (Wald test, individual *P* values > 0.05, FDR corrected). In both genotypes, among over 2.8 million gene abundances in 2,545 genes, at least the 50% of the genes was included in pathways within the ‘Biofertiliazion’, ‘Colonizing plant system’, ‘Biocontrol’, ‘Competitive exclusion’, ‘Bioremediation’ and ‘Phytohormone and Plant Signaling’ categories [Fig. 6A, B; Data Set S4 at https://zenodo.org/records/17580416]. Regardless of the host genotype, the PGPTs associated with *A. calcoaceticus* metagenomic reads were mostly represented by functions related to organic acid metabolism and ROS scavenging, involved in phosphate assimilation and biocontrol activities, respectively. Despite the functional conservation among tomato genotypes [Fig. 6; Data Set S4 at https://zenodo.org/records/17580416], ∼10% PGPTs were uniquely identified in ‘Ab’ and ‘SV’ Acinetobacter MAG profiles, representing a possible “footprint” of host selection on the rhizosphere microbiota [Fig. S7; Data Set S4 at https://zenodo.org/records/17580416]. We next mined for ARGs into the Acinetobacter MAG: intriguingly we failed to identify significant “hits” in multiple databases, with the exception of the identification of an antibiotic efflux pump *AbaQ* previously implicated in Quinolone resistance (60).

**Figure 6:**
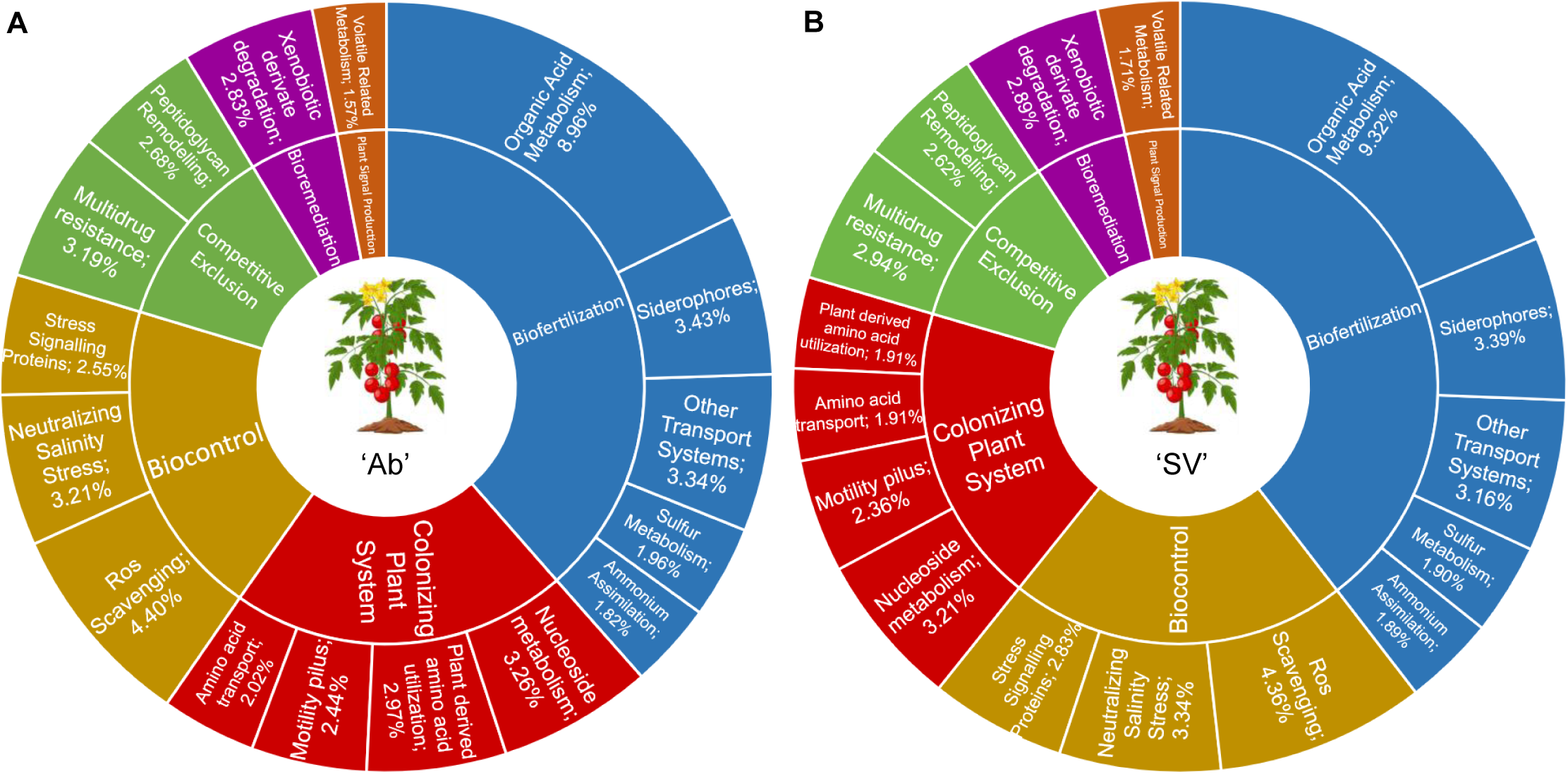
Average proportions of the most representative gene abundances associated with PGPT categories at level four, based on metagenomic reads assigned to *Acinetobacter calcoaceticus* in (**A**) ‘Ab’ and (**B**) ‘SV’ genotypes.

Finally, to gain further insights into the functional potential of the tomato microbiota we expanded our investigation to eight additional MAGs [Table S5]. With the caveat that these MAGs represent taxa whose abundance is at least an order of magnitude less than Acinetobacter, their enriched functions allowed us to identify three main categories, i.e., Metabolism, Environmental Information and Genetic Information processing, displaying a ‘microhabitat’ effect in both genotypes [Wald test, individual *P* values < 0.05, FDR corrected; Data Set S5 at https://zenodo.org/records/17580416].

## DISCUSSION & LIMITATIONS

Similar to what previously reported, the tomato plants grown in this “lab-in-the-field” hosted a bacterial community whose richness and diversity decrease from the unplanted soil to the inner root tissues and communities are dominated by relatively few taxa (18).

A striking feature of our investigation is the dominance of an individual *Acinetobacter sp*. of the rhizosphere and root profiles. This dominance has previously been reported for tomato growing both in glasshouse (61) and field trials (62), and Acinetobacter can be isolated from tomato plants (63). This motivated us to discern origin and functional significance of the observed dominance. For instance, it is known that exogenous factors may act as a reservoir for the tomato microbiota, as reported for rain and the phyllosphere microbiota (64). However, we found no evidence for a passive flow-through of Acinetobacter from the microbiota retrieved from different farm machinery. An alternative scenario is represented by the opportunistic nature of Acinetobacter colonisation, as previously reported for *Ralstonia*-infected tomato plants (65). However, no evidence for disease establishment was reported for the investigated field. This left us with two plausible explanations. The first one is a seed-transmission of microbes to mature plants, a scenario congruent with what reported for beneficial bacteria inhabiting tomato inner tissues (66). The second explanation is represented by an enrichment from the surrounding soil biota of Acinetobacter. This latter explanation would be congruent with what we previously reported for field-grown tomato plants in the north of Italy (19). Time-course experiments, profiling the development of the tomato microbiota from seeds to transplanted seedlings to established crops will be required to validate either [or both] scenarios.

As Acinetobacter dominance defines both below-ground tomato microhabitats, we reasoned that a comparative metagenomic analysis of the rhizosphere and unplanted soil specimens would have provided us with insights into the biological significance of the bacterial enrichment. Consistently, the taxonomic composition of the obtained metagenomes reflected a) the substantial congruence between the data obtained with different sequencing strategies and b) a bacterial dominance. This motivated us to infer functional significance using a two-pronged approach. First, we looked at the differential enrichment of a curated dataset of plant-growth promoting functions (51), and we observed how mineral mobilisation, including iron, phosphorus and sulphur emerged as the dominant functions enriched in the tomato rhizosphere in a genotype-dependent manner. Other striking enrichments were represented by the adaptation to salinity and osmotic stresses as well as detoxification of reactive oxygen species, as those processes are intertwined (67). As salinisation is a developing threat for agricultural production in Mediterranean environments (68), these enrichments can be interpreted as evidence of the impact of climatic modifications on the tomato microbiota.

Taken together, these observations are congruent with the recruitment of a versatile plant growth-microbe at the tomato root-soil interface, fitting the profile of soil-borne Acinetobacter (69). Yet, although beneficial for their host, this recruitment may be a Janus-faced one. For instance, the genus Acinetobacter has been of clinical interest for decades (70), with *Acinetobacter baumanni* considered a paradigmatic examplefor its vast multidrug resistance capabilities (71). Alongside this established case study, other members of the genus are gaining centre-stage for their antimicrobial resistance (72, 73). For instance, naturally abundant *Acinetobacter sp*. in plant food chains may act as a vehicle for ARGs distribution in clinical settings (74), although ‘non-human’ strains tend to have fewer antibiotic resistance genes, mostly intrinsic in nature (75). Consistently, we failed to obtain evidence that the betalactamase dominance identified in the community resistome links taxonomic composition with ARG distribution in the tomato microbiota, as previously reported across soil habitats (76). Despite the genus Acinetobacter as a whole is recognised as a natural reservoir of betalactamase (77, 78) our results concur with a recent investigation of non-clinical isolates of *A. baumanni* from grasslands displaying a limited betalactamase distribution [coupled though with the presence of *AbaQ*] (79). Consequently, if ARGs confer an adaptive advantage to Acinetobacter colonisation, this appears a) habitat-specific, as we did not observe any enrichment for this bacterium in the phyllosphere, and b) decoupled from the ‘community resistome’, that are ARGs coded by other members of the community. This result is aligned with previous observations (80) indicating that the soil resistome appears driven by abiotic factors, including temperature and precipitation, as opposed to the phyllosphere resistome responding to inter-microbial competition.

Our work suggests a scenario whereby the rhizosphere enrichment of a plant beneficial bacterium, is mediated, at least in part, by ARGs. Though drought tolerance is often associated to members of the phylum Actinobacteria (81), a recent investigation identified an Acinetobacter from arid soils among bacterial capable of alleviating tomato drought stress (82). It is therefore tempting to speculate that the observed Acinetobacter enrichment may represent a “plant’s attempt” at future-proofing the microbiota for better adaptation to climatic modifications. This is reminiscent the Pseudomonas phylotypes “bloom” occurred in a host genotype-independent manner at the in the sorghum microbiota (83) : dominant phylotypes, unlike “hubs”, respond rapidly to substrate- and/or environmental-driven fluctuations and may establish with poor connectivity with the rest of the microbiota. With increased knowledge of the host genetic determinants of the tomato microbiota (15), beneficial microbial enrichment could be soon integrated into plant breeding strategies (3). In doing so, it will however be important to consider potential trade-offs between microbial configurations and tomato performance, as highlighted by both ecological (84) and molecular (85) investigations. This may include the characterisation of isolates of the identified Acinetobacter, complementing surveys envisioned for the genera (86) and/or for other plant growth-promoting bacteria (87). In turn, this will provide scientists and practitioners with a better understanding of the ‘One Health impact’ of the deliberate manipulation of individual members of the plant microbiota.

A limitation of the study is that we report the results of an individual year of experimentation from an individual farm. As plant development is known to fine-tune the recruitment cues of the microbiota, as previously reported by ourselves (19) and other groups (88), the individual experiment prevented us from inferring the contribution of season-to-season variation in shaping the tomato microbiota. Further experimentations, across growing seasons and farms will elucidate the deterministic nature of microhabitat and host genetic factors in defining composition and function of the microbiota thriving in association with field-grown plants.

## DATA AVAILABILITY

The genomic sequences reported in this study are deposited in the European Nucleotide Archive (ENA), project PRJEB80388. The codes to reproduce data analyses are available at https://github.com/BulgarelliD-Lab/CIRCLES_Tomato

## AUTHOR CONTRIBUTIONS

SR, amplicon data analysis; AM, metagenomic data analysis; RAT, LP and PDC, sampling; SA, AC, HNN sequencing libraries preparation; JM, PH, HNN, RH and CCL, sequencing data generation and inspection; CA and PT, Acinetobacter phylogenetic analysis; FDM, FMA and PL, ARG data analysis; GB and PLM network analysis; CS, data archiving and preparation for dissemination; SR, AM and DB, draft writing; all co-authors reviewing and editing; SR, AM, JA and DB, conceptualization; PDC,, PLM, FMA, PW and DB, funding acquisition.

## ACKNOWLWDGEMENTS

This work was supported by the Horizon 2020 Framework Program Innovation Action “CIRCLES” (European Commission grant agreement 818290) awarded to the University of Dundee, the University of Bologna, Luxembourg Centre for Systems Biomedicine, DTU Food and MS Biotech. CA is supported by a UKRI BBSRC eastBIO PhD studentship (project 2734186).

The authors are thankful to the farmers who provided us with access to their fields. We thank Elena Del Rio and Jen Gallagher [University of Dundee] for their administrative and technical support, respectively, in delivering the project.

