## Supplementary Information for "Acinetobacter enrichment shapes composition and function of the bacterial microbiota of field-grown tomato plants"

*Running Head:* *The bacterial microbiota of field-grown tomato plants*

Keywords: food-chain microbiota, tomato, metagenomics, AMR, One Health

Present address:

Senga Robertson: School of Health Science, University of Dundee, Dundee, UK

Rodrigo Alegria Terrazas: Rothamsted Research, Harpenden, UK

### **SUPPLEMENTARY TABLES**

### **Table S1. Soils main chemical and physical properties**

|  | ‘Ab’ soil | ‘SV’ soil |
| --- | --- | --- |
| Sand (%) | 20 | 19 |
| Silt (%) | 59 | 58 |
| Clay (%) | 21 | 23 |
| pH^1^ | 7.8 | 8.1 |
| Organic Carbon^2^ (g/Kg) | 16.8 | 14 |
| Total nitrogen^3^ (g/Kg) | 2.9 | 3.3 |
| Available phosphorous^4^ (mg/Kg) | 130 | 148 |
| C/N ratio | 5.8 | 4.3 |

^1^water extract; ^2^Walkey-Black determination; ^3^Kjeldhal determination; ^4^Olsen determination

### **Table S2. Co-occurrence networks: topology and centrality genera among microhabitats in both genotypes**

|  | Nodes/Edges | Degree  centrality | Betweenness centrality | Closeness  centrality |
| --- | --- | --- | --- | --- |
| Bulk | 82/282 | *Ellin517*  *Sphingomonas* | *Sphingomonas* | *Sphingomonas* |
| Leaves | 16/48 | *-* | *Allorhizobium-Neorhizobium-Pararhizobium-Rhizobium* | *Allorhizobium-Neorhizobium-Pararhizobium-Rhizobium*  *Pantoea*  *Massilia* |
| Rhizosphere | 23/50 | *Limnohabitans Pseudomonas*  *Terribacillus* | *Limnohabitans Pseudomonas*  *Terribacillus* | *Terrabacillus* |
| Roots | 19/24 | *Lysinibacillus* | *-* | *Lysinibacillus* |

### **Table S3. Metagenomic assembly information**

|  | ‘Ab’ samples | ‘SV’ samples |
| --- | --- | --- |
| Assembly length (bp) | 787,670,201 | 834,028,659 |
| Number of contigs | 1,262,321 | 1,112,553 |
| Largest contig (bp) | 58,515 | 71,853 |
| *N50* (bp) | 673 | 909 |
| *L50* (bp) | 341,714 | 253,927 |

**Table S4. Proportion of a metagenomic-assembled genome (MAG) identified as *Acinetobacter calcoaceticus* across the host-filtered metagenomic sequences of the 37 tomato samples.**

| **Sample** | **Mapping (%)** | **Microhabitat** | **Genotype** |
| --- | --- | --- | --- |
| T1_B2 | 0.92% | Rhizosphere | 'Ab' |
| T1_D2 | 0.94% | Rhizosphere | 'Ab' |
| T1_E2 | 2.19% | Rhizosphere | 'Ab' |
| T2_A2 | 5.07% | Rhizosphere | 'Ab' |
| T2_B2 | 10.43% | Rhizosphere | 'Ab' |
| T2_C2 | 0.39% | Rhizosphere | 'Ab' |
| T2_D2 | 1.32% | Rhizosphere | 'Ab' |
| T2_E2 | 2.84% | Rhizosphere | 'Ab' |
| T2_S | 0.07% | Bulk | Unplanted |
| T3_C2 | 3.20% | Rhizosphere | 'Ab' |
| T3_E2 | 14.49% | Rhizosphere | 'Ab' |
| T3_S1 | 0.86% | Bulk | Unplanted |
| T3_S1 | 0.49% | Bulk | Unplanted |
| T4_A2 | 1.12% | Rhizosphere | 'Ab' |
| T4_B2 | 10.39% | Rhizosphere | 'Ab' |
| T4_C2 | 2.87% | Rhizosphere | 'Ab' |
| T4_D2 | 5.39% | Rhizosphere | 'Ab' |
| T5_B2 | 2.74% | Rhizosphere | 'Ab' |
| T5_C2 | 4.94% | Rhizosphere | 'Ab' |
| T5_D2 | 4.61% | Rhizosphere | 'Ab' |
| T10_A2 | 3.25% | Rhizosphere | 'SV' |
| T10_C2 | 3.09% | Rhizosphere | 'SV' |
| T10_D2 | 0.57% | Rhizosphere | 'SV' |
| T10_E2 | 1.58% | Rhizosphere | 'SV' |
| T6_B2 | 5.26% | Rhizosphere | 'SV' |
| T6_C2 | 2.76% | Rhizosphere | 'SV' |
| T6_D2 | 3.55% | Rhizosphere | 'SV' |
| T6_E2 | 4.64% | Rhizosphere | 'SV' |
| T7_B2 | 6.30% | Rhizosphere | 'SV' |
| T7_C2 | 5.24% | Rhizosphere | 'SV' |
| T7_D2 | 0.05% | Rhizosphere | 'SV' |
| T7_E2 | 4.79% | Rhizosphere | 'SV' |
| T7_S | 0.91% | Bulk | Unplanted |
| T8_S1 | 2.02% | Bulk | Unplanted |
| T9_B2 | 4.79% | Rhizosphere | 'SV' |
| T9_D2 | 0.88% | Rhizosphere | 'SV' |
| T9_E2 | 1.96% | Rhizosphere | 'SV' |

**Table S5. General statistics of the MAGs selected according to the completeness and contamination parameters.**

| **Phylum** | **Genus or Species** | **Completeness (%)** | **Contamination (%)** |
| --- | --- | --- | --- |
| Thermoprotetota | TA-21 | 78.16 | 2.91 |
| Bacteroidota | *Flavobacterium* | 73.98 | 3.46 |
| Proteobacteria | *Acinetobacter calcoaceticus* | 72.15 | 8.87 |
| Bacteroidota | *Sphingobacterium* sp002472835 | 52.99 | 4.05 |
| Bacteroidota | *Chryseobacterium* | 49.72 | 1.63 |
| Bacteroidota | *Pedobacter ginsengisoli* | 41.64 | 2.1 |
| Bacteroidota | *Flavobacterium* | 78.45 | 10.38 |
| Bacteroidota | *Flavobacterium* | 62.93 | 0.86 |
| Bacteroidota | *Flavobacterium* | 49.23 | 13.79 |

### **SUPPLEMENTARY FIGURES**

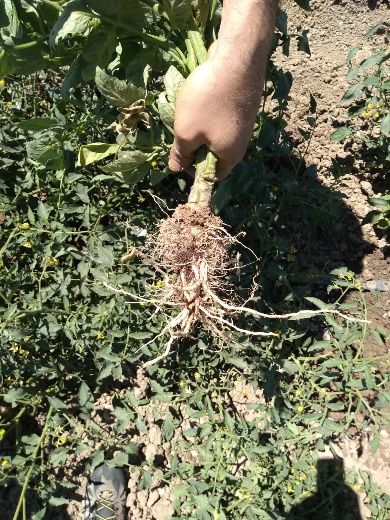

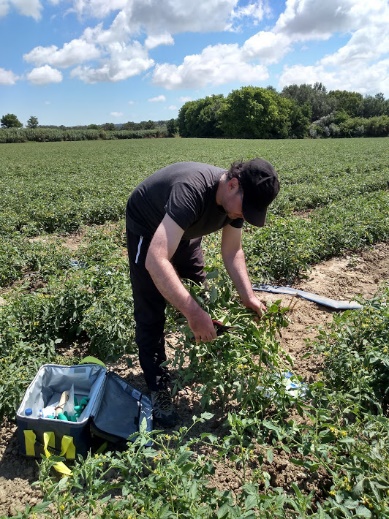

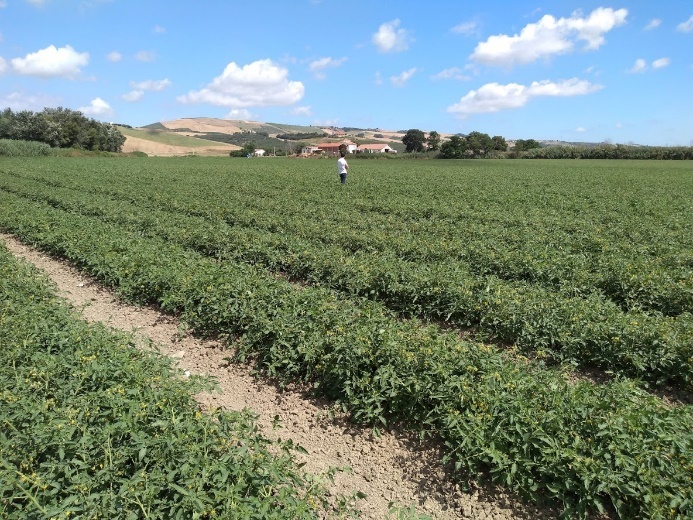

### **Figure S1. Representative pictures of the fields and plants at the time of sampling. *The person depicted in the photographs is one of the co-authors of the manuscript.***

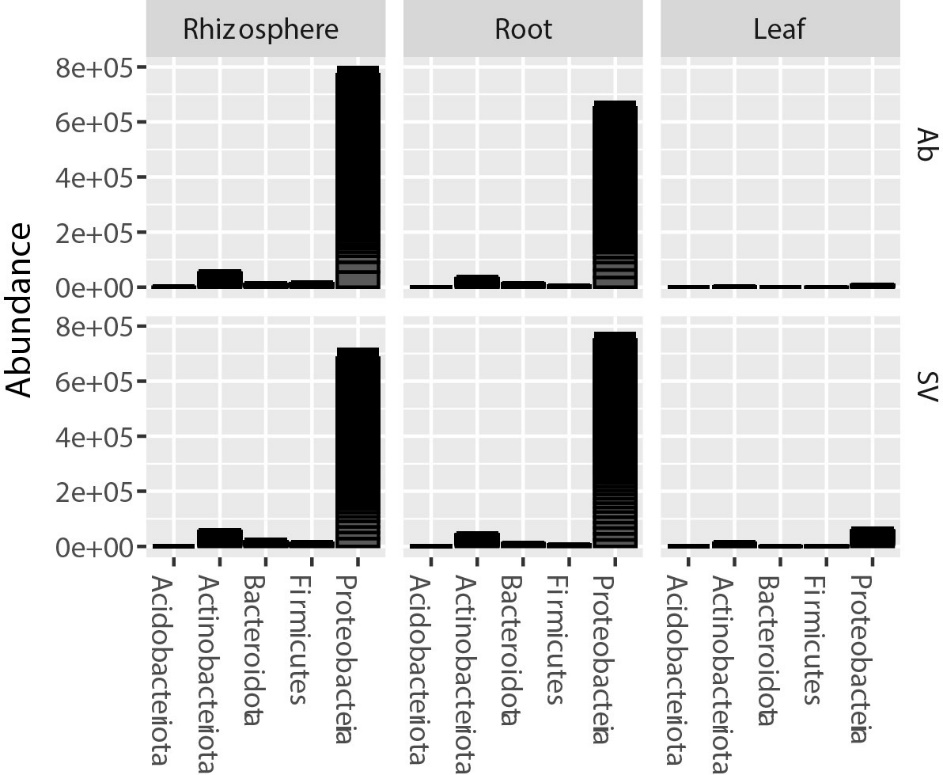

### **Figure S2. Taxonomic composition of the tomato microhabitats.**

Cumulative relative abundance, expressed as count per millions, of bacterial ASVs identified in this study grouped at phylum level in the indicated microhabitats. In each “tower”, individual blocks depict biological replicates for given ASVs

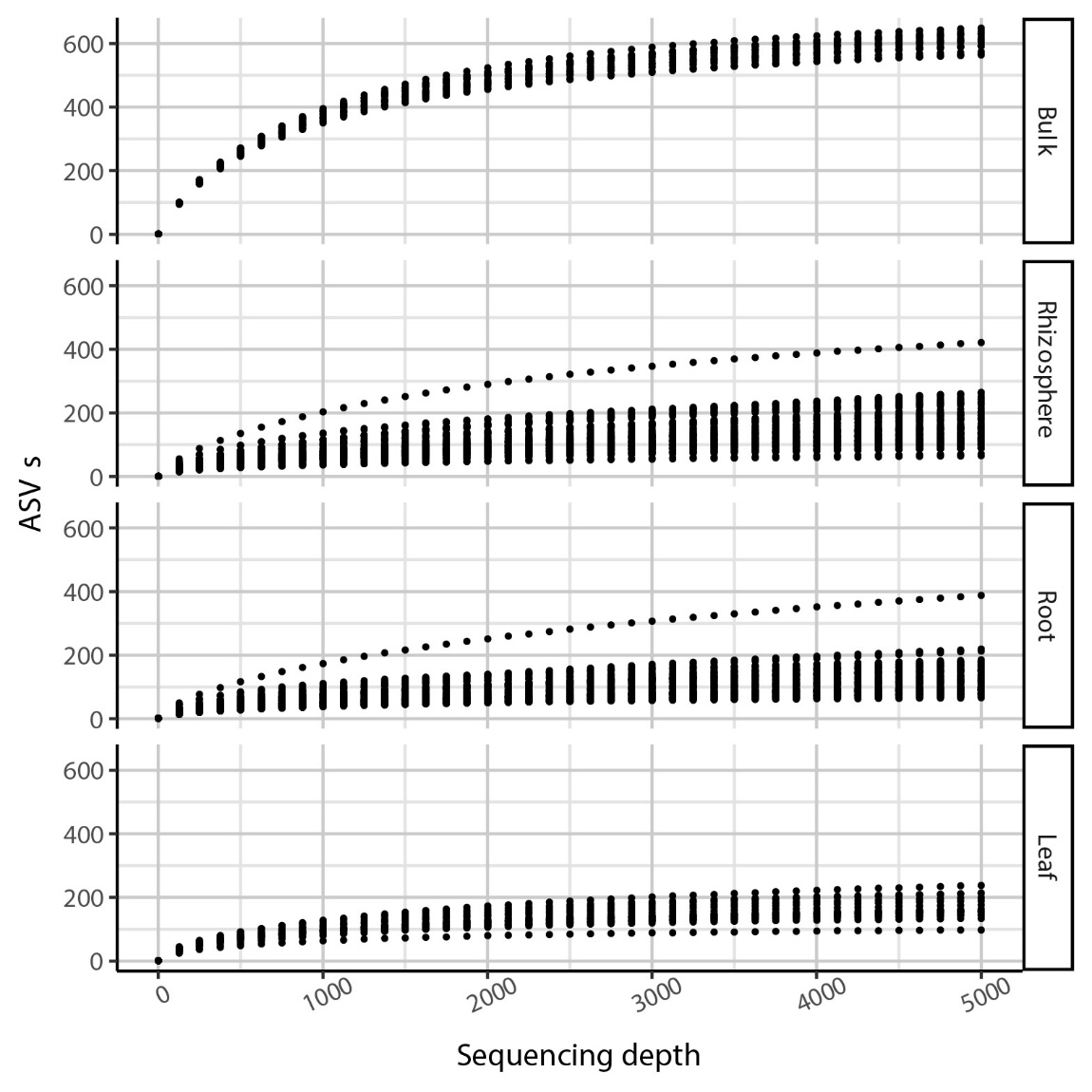

### **Figure S3. Rarefaction curves for the investigated microhabitats.**

For each microhabitat, individual dots depict number of identified ASVs for a given biological replicate collated and displayed at 125-read sequencing depth interval.

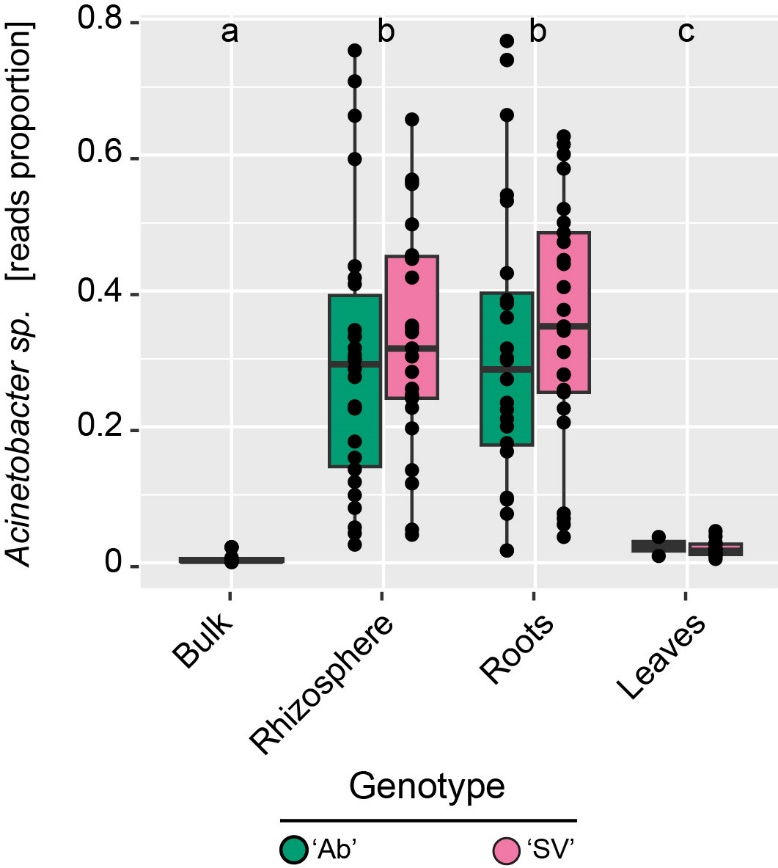

### **Figure S4. *Acinetobacter sp.* dominates root and rhizosphere microhabitats.**

Proportion of sequencing reads [ratio Acinetobacter reads/total reads per individual sample] across microhabitat and genotypes. The upper and lower edges of the box plots represent the upper and lower quartiles, respectively. The bold line within the box denotes the median, individual shapes depict measurements of individual biological replicates/genotypes for a given microhabitat. Different letters denote statistically significant differences between microhabitat means by Kruskal–Wallis non-parametric analysis of variance followed by Dunn’s post-hoc test (Individual *P values* < 0.001; BH corrected).

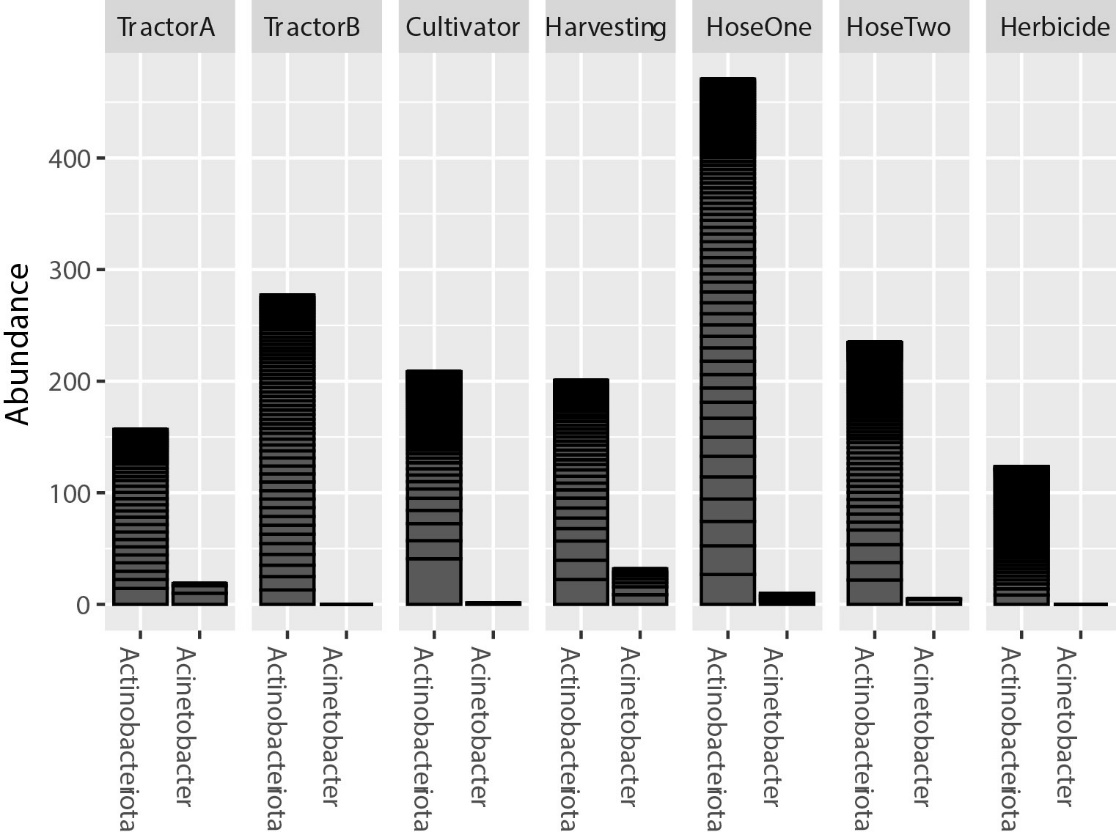

### **Figure S5. Actinobacteriota dominate the farm machinery microbiota**

Cumulative relative abundance, expressed as count per millions, of bacterial ASVs assigned to Actinobacteriota or Acinetobacter *sp*. in the indicated farm machinery. In each “tower”, individual blocks depict biological replicates for given ASVs. ‘Herbicide’ denotes herbicide sprayer.

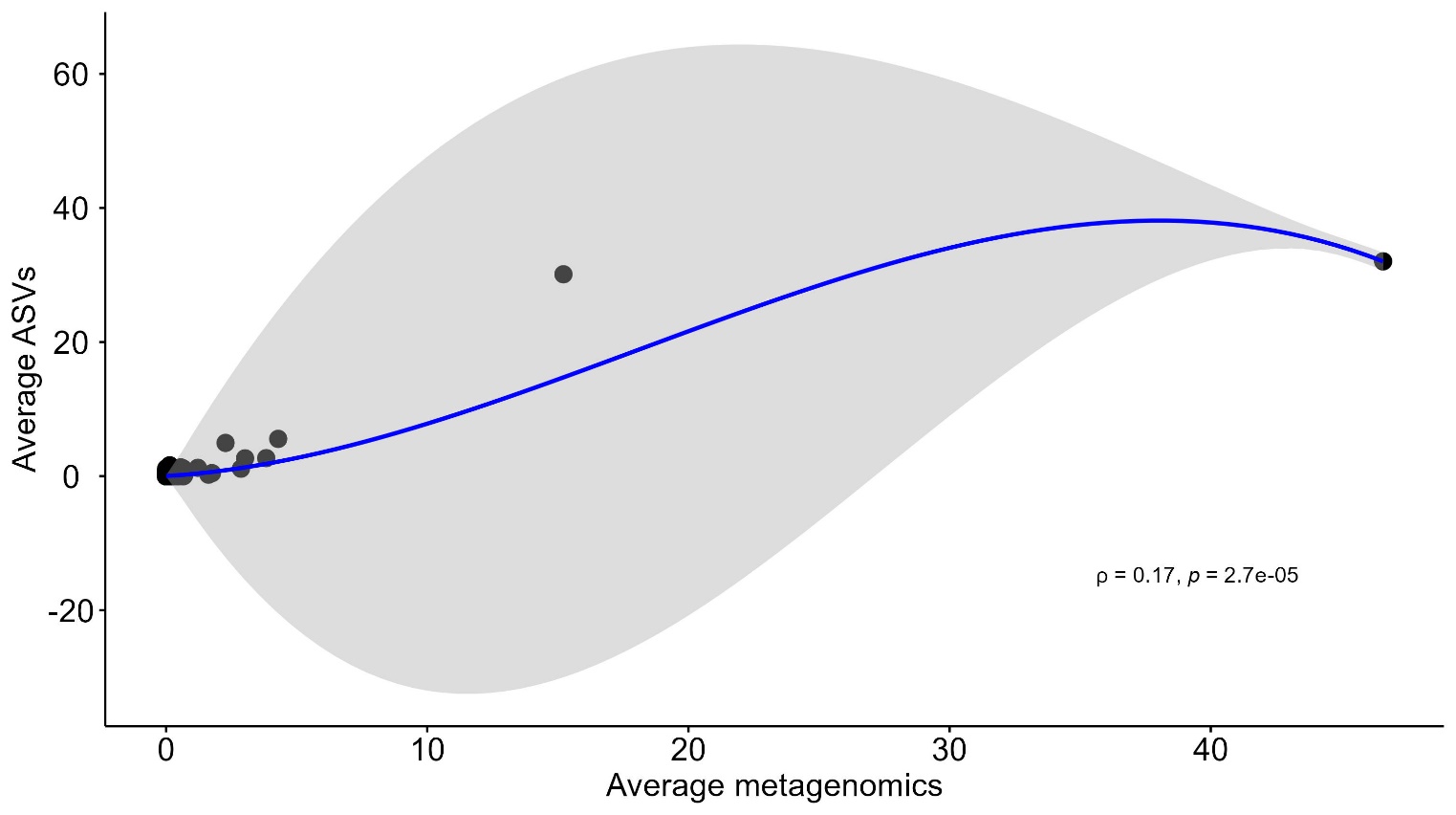

### **Figure S6. Spearman’s rank correlation between taxonomic composition amplicon sequencing and metagenomics.**

Scatter plot depicting the statistically significative (*P* value < 0.05) Spearman’s rank correlation between the average relative abundance of the metagenomic and ASVs reads at the family taxonomic level in the x and y axis, respectively. The area coloured in grey represents the confidence interval, while the blue line the local regression (LOESS) curve.

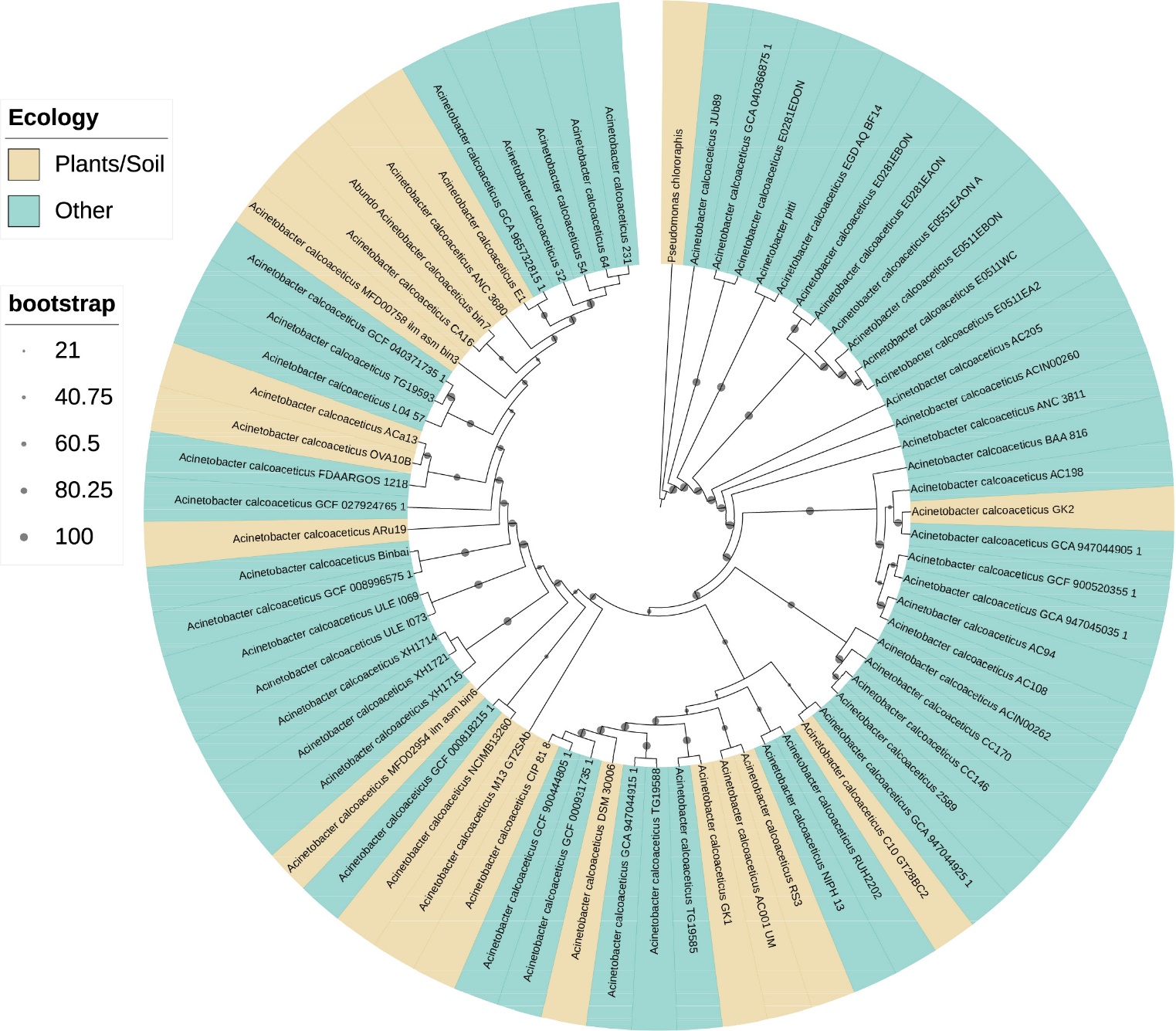

### **Figure S7. Phylogenetic relatedness of the Acinetobacter MAG and other *Acinetobacter calcoaceticus* genomes.**

Whole-genome phylogenetic tree of individual Acinetobacter genomes constructed with 100 bootstrap iterations. The colour-coding depicts strain’s ecological niche; ‘other’ include clinical, environmental or strains for which the isolation source is not reported in NCBI. The arrowhead depicts the tree position of the Acinetobacter MAG.

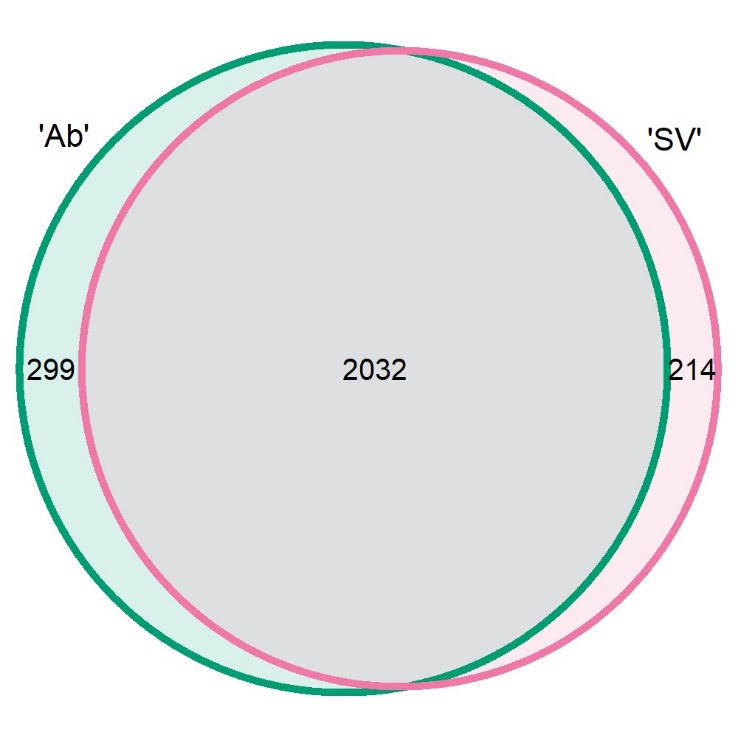

### **Figure S8. Unique and shared PGPTs between ‘Ab’ and ‘SV’ *Acinetobacter calcoaceticus* metagenomic reads.**

Venn diagram showing the distribution of PGPTs among genotypes. Values within each circle represent the number of PGPTs, coloured according to the host genotype. The overlapping area indicates PGPTs shared between the genotypes.
